# Scirpy: A Scanpy extension for analyzing single-cell T-cell receptor sequencing data

**DOI:** 10.1101/2020.04.10.035865

**Authors:** Gregor Sturm, Tamas Szabo, Georgios Fotakis, Marlene Haider, Dietmar Rieder, Zlatko Trajanoski, Francesca Finotello

## Abstract

**Summary:** Advances in single-cell technologies have enabled the investigation of T cell phenotypes and repertoires at unprecedented resolution and scale. Bioinformatic methods for the efficient analysis of these large-scale datasets are instrumental for advancing our understanding of adaptive immune responses in cancer, but also in infectious diseases like COVID-19. However, while well-established solutions are accessible for the processing of single-cell transcriptomes, no streamlined pipelines are available for the comprehensive characterization of T cell receptors. Here we propose *Scirpy*, a scalable Python toolkit that provides simplified access to the analysis and visualization of immune repertoires from single cells and seamless integration with transcriptomic data.

**Availability and implementation:** *Scirpy* source code and documentation are available at https://github.com/icbi-lab/scirpy.

## Introduction

B and T lymphocytes are equipped with a vast repertoire of immune cell receptors that can recognize a wealth of different antigens. High-throughput sequencing technologies have enabled the study of these immune repertoires at unprecedented resolution (Hackl *et al.*, 2016; Finotello *et al.*, 2019) and are advancing our understanding of adaptive immune responses in cancer (Valpione *et al.*, 2020), as well as in autoimmune (Hanson *et al.*, 2020) and infectious (Schober *et al.*, 2020) diseases, including COVID-19.

Novel single-cell sequencing technologies now allow the joint profiling of transcriptomes and T cell receptors (TCRs) in single cells. However, while the study of single-cell transcriptomes is facilitated by tools like Seurat (Butler *et al.*, 2018) and Scanpy (Wolf *et al.*, 2018), the bioinformatic analysis of paired α and β TCR chains is still in its infancy. Several methods to perform specific analytical tasks have been proposed (**Supplementary Table 1**), but the comprehensive characterization of TCR diversity from single cells is still hampered by the lack of standardized and ready-to-use computational pipelines.

Here, we present *Scirpy* (**s**ingle-**c**ell **i**mmune **r**epertoires in **Py**thon), a Python-based Scanpy extension that provides simplified access to various computational modules for the analysis and visualization of immune repertoires from single cells. Due to its tight integration with Scanpy, *Scirpy* allows the combination with scRNA-seq transcriptomic data to comprehensively characterize the phenotype and TCR of single T cells.

## The *Scirpy* package

*Scirpy* integrates different bioinformatic methods for importing, analyzing, and visualizing single-cell TCR-sequencing data (Fig. 1). TCR data can be loaded from CellRanger (10x Genomics) and TraCeR (Stubbington *et al.*, 2016) outputs, thus allowing the analysis of both 10x Genomics and Smart-seq2 data, respectively. The *AnnData* data structure provided by Scanpy is used to store matched TCR information and transcriptomic profiles.

**Figure 1.**
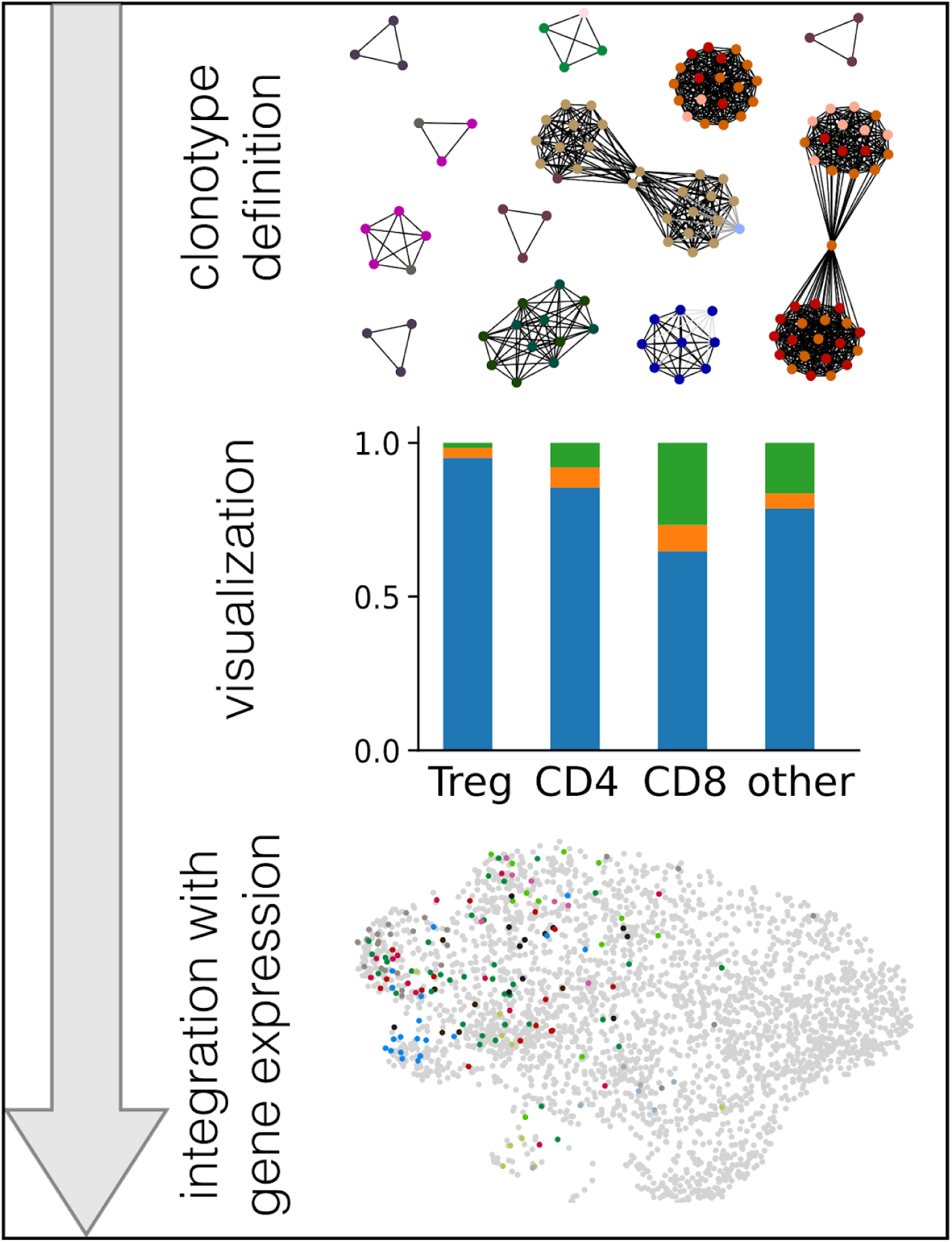
The *Scirpy* workflow. After defining clonotypes via CDR3-sequence similarity, *scirpy* offers a wide range of visualization options to explore clonotype expansion, abundance, and VDJ gene usage. Finally, clonotype information can be integrated with transcriptomic data, leveraging the scanpy workflow. Top panel: Exemplary clonotype network. Each node represents a cell, colored by sample. Edges connect cells belonging to the same clonotype. Middle panel: Clonal expansion of different T cell subsets visualized as bar chart. The bars colored in blue, orange, and green represent the fractions of cells belonging to clonotypes with one, two or more than two cells, respectively. Lower panel: UMAP embedding based on gene expression. Colored dots represent the cells belonging to the most abundant clonotypes.

*Scirpy* uses a flexible TCR model supporting up to two α and β chains per cell, allowing the identification of *dual-TCR T cells* (Schuldt and Binstadt, 2019) (see **Supplementary Note 1**). It also flags cells with more than two chains, which potentially represent doublets (**Supplementary Fig. 1**) and may be discarded from downstream analyses. *Scirpy* defines clonotypes based on the amino-acid or nucleotide sequence of the TCR complementarity-determining region 3 (CDR3). The user can choose between defining clonotypes based on sequence identity or similarity. The second approach, inspired by TCRdist (Dash *et al.*, 2017), leverages the Parasail library (Daily, 2016) to compute pairwise sequence alignments and identify clusters of T cell clonotypes that likely recognize the same antigens. For building clonotype networks, *Scirpy* makes use of the sparse-matrix implementation from the *scipy* package (Virtanen *et al.*, 2020), ensuring scalability to hundreds of thousands of cells.

*Scirpy* offers a wide range of tools and visualization options that we demonstrate in the “case study” section. It allows inspecting TCR chain configurations **(Supplementary Fig. 1)**, and exploring the abundance, diversity, and expansion of clonotypes across samples, patients, or cell-type clusters derived from transcriptomics data (**Supplementary Fig. 2** and **3**). Relationships between clonotypes can be investigated with a graph-based approach **(Supplementary Figure 4)**, in addition to spectratype plots that visualize CDR3 sequence length distribution, and VDJ-usage plots **(Supplementary Figure 5)**. Finally, TCR information can be integrated with transcriptomic data, for instance by overlaying Uniform Manifold Approximation and Projection (UMAP) plots (**Supplementary Figure 3**). A detailed tutorial guiding through a typical analysis workflow is available at https://icbi-lab.github.io/scirpy/tutorials.html.

## Case study: re-analysis of 140k single T cells

To demonstrate the applicability to a real-world scenario, we re-analyzed a recent single-cell dataset of ~140k T cells (Wu *et al.*, 2020). Single T cells were isolated from tumours, normal adjacent tissue, and peripheral blood of 14 patients with four different cancer types, and subjected to single-cell RNA and TCR sequencing with the 10x technology. Consistently with the original results, we found that the majority of clonotypes were singletons and only 8-19% of patients’ clonotypes were clonally expanded **(Supplementary Fig. 2)**. Our results further confirm that CD8+ effector, effector memory, and tissue resident T cells comprised a large fraction of clonotypes that were expanded in both the tumor and normal tissue, while CD4+ T cells consisted mostly of singletons **(Supplementary Fig 3)**. Moreover, leveraging Scirpy’s capability to define clonotypes based on sequence-similarity rather than identity, we identified clusters of CDR3 amino-acid sequences indicating convergent TCR evolution **(Supplementary Fig. 4)**.

The analysis ran in 12 minutes on a single core of an Intel E5-2699A v4, 2.4 GHz CPU when defining clonotypes based on sequence identity, and in 35 minutes on 44 cores when using pairwise sequence alignment. A jupyter notebook to reproduce this case study is available at: https://icbi-lab.github.io/scirpy-paper/wu2020.html.

## Conclusions

*Scirpy* is a versatile tool to analyze single-cell TCR-sequencing data that enables seamless integration with the Scanpy toolkit, the *de facto* standard for analyzing single-cell data in Python. *Scirpy* is highly scalable to big scRNA-seq data and, thus, allows the joint characterization of phenotypes and immune cell receptors in hundreds of thousands of T cells. An extension of *Scirpy* to characterize γδ TCR and B cell receptor (BCR) repertoires is planned for the next release.

## Supporting information

figure1 (high resolution)

