## supplementary information for "Scirpy: A Scanpy extension for analyzing single-cell T-cell receptor sequencing data"

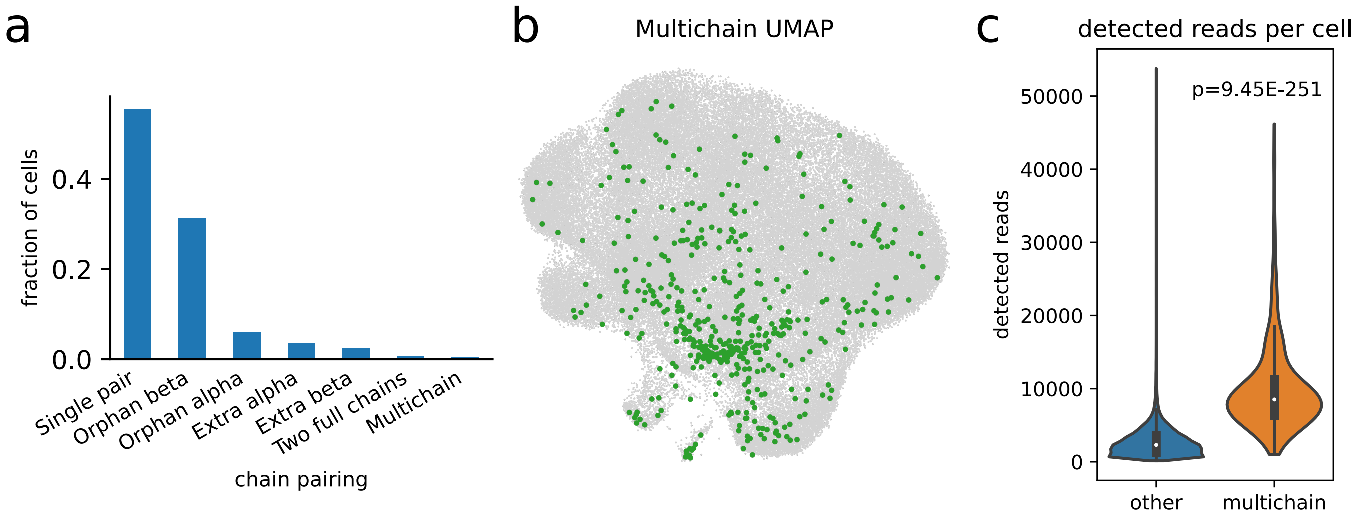


**Supplementary Figure 1: Scirpy flags cells with more than two pairs of α- and β-TCR-chains as "multichain" cells.** **(a)** The chart delineates the fraction of cells with a certain receptor configuration. Orphan alpha and beta designates cells where only a single α- or β-chain sequence could be recovered, respectively; extra refers to cells having an extra chain in addition to a valid pair of α- or β-receptor chains. **(b)** UMAP plot of 96,000 cells from Wu *et al.* [1] with at least one detected CDR3 sequence with multichain-cells (n=474) highlighted in green. **(c)** Comparison of detected reads per cell in multichain-cells and other cells. Multichain cells comprised significantly more reads per cell ($p = 9.45 \times 10^{-251}$, Wilcoxon-Mann-Whitney-test), supporting the hypothesis that (most of) multichain cells are technical artifacts arising from cell-multiplets [2].

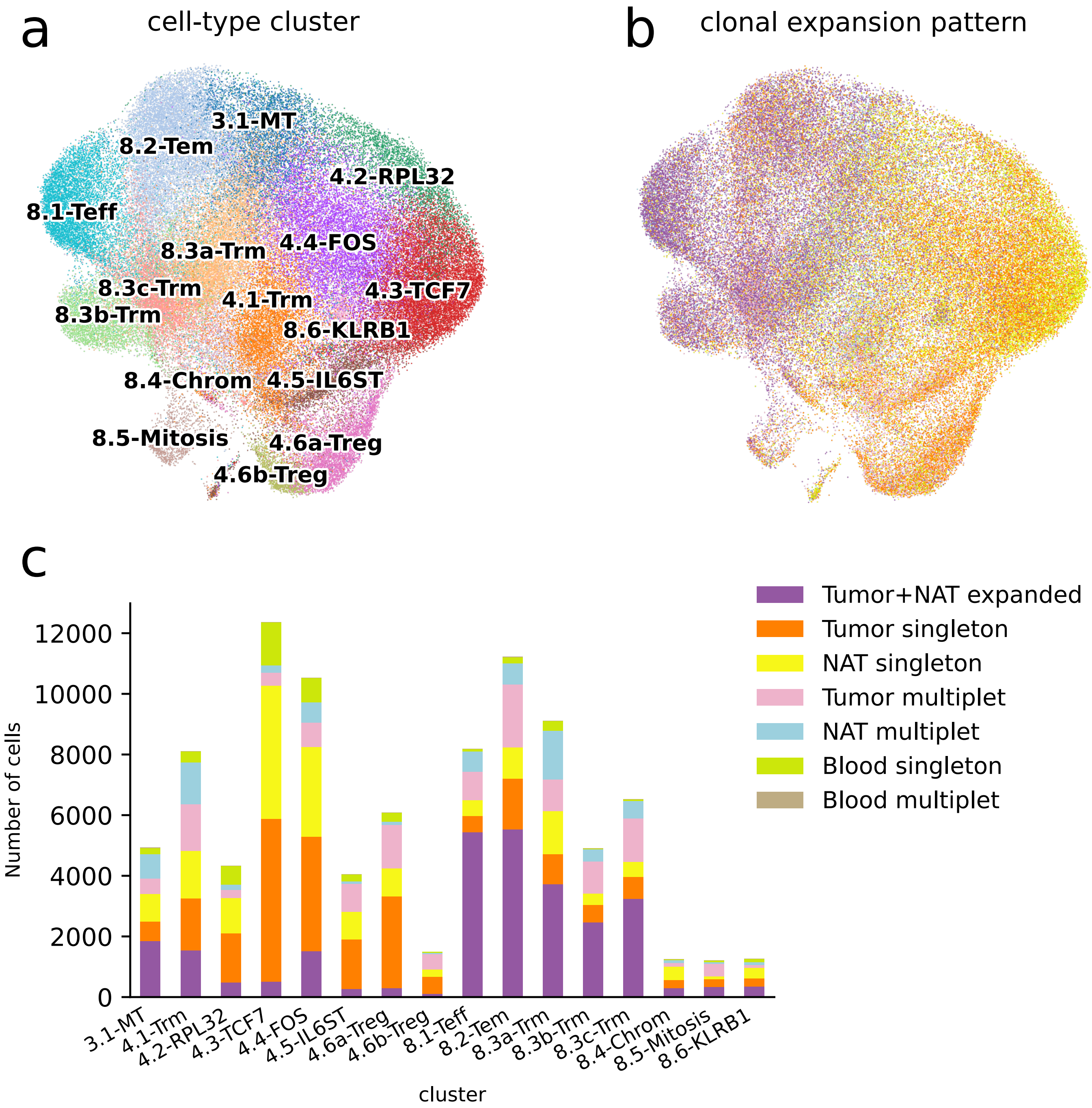


**Supplementary Figure 2: Differential tissue expansion patterns by T-cell clusters. (a)** UMAP plot of 96,000 cells from Wu *et al.* [1] with at least one detected CDR3 region, colored according to the cell-type clusters described in the original study. **(b, c)** Distribution of tissue expansion patterns by clustered T-cell subpopulations visualized as (b) UMAP plot and (c) bar chart. While CD8+ effector, effector memory, and tissue-resident T-cells tended to be expanded in both tumor and normal adjacent tissue (NAT), CD4+ T-cell clusters primarily comprised singleton clonotypes.

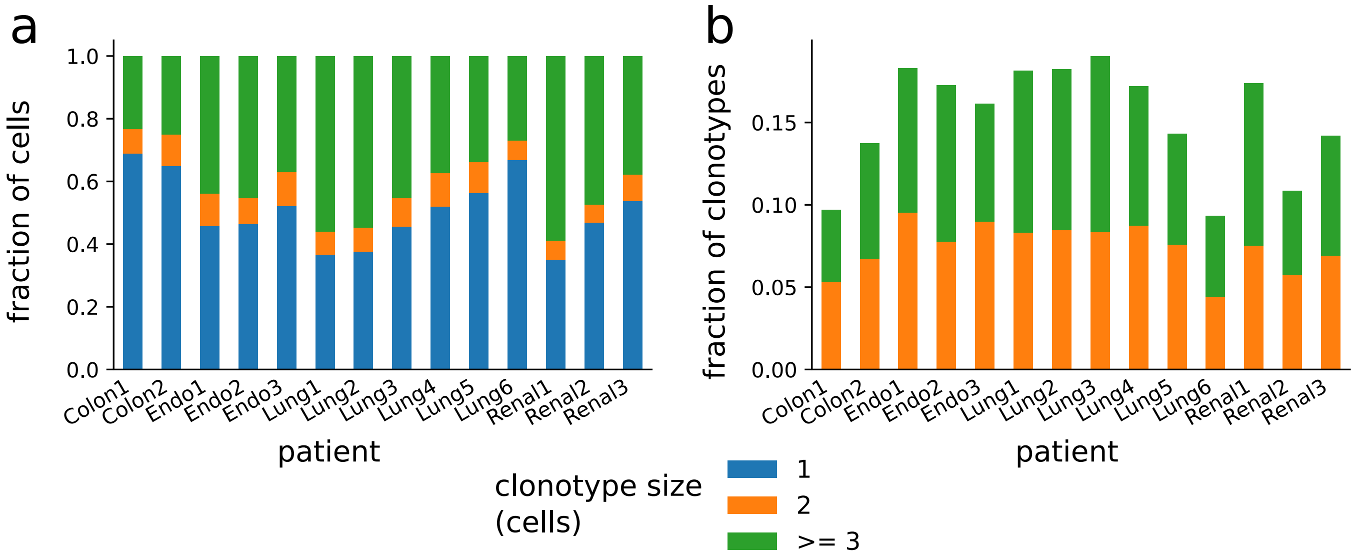


**Supplementary Figure 3: T cell clonal expansion per patient.** **(a)** Fraction of cells belonging to clonotype singletons (blue), clonotypes with two cells (orange), or clonotypes with more than two cells (green). **(b)** Fraction of clonotypes comprising two, or more than two cells, respectively. Overall, between 8% and 19% of clonotypes were expanded, depending on the patient.

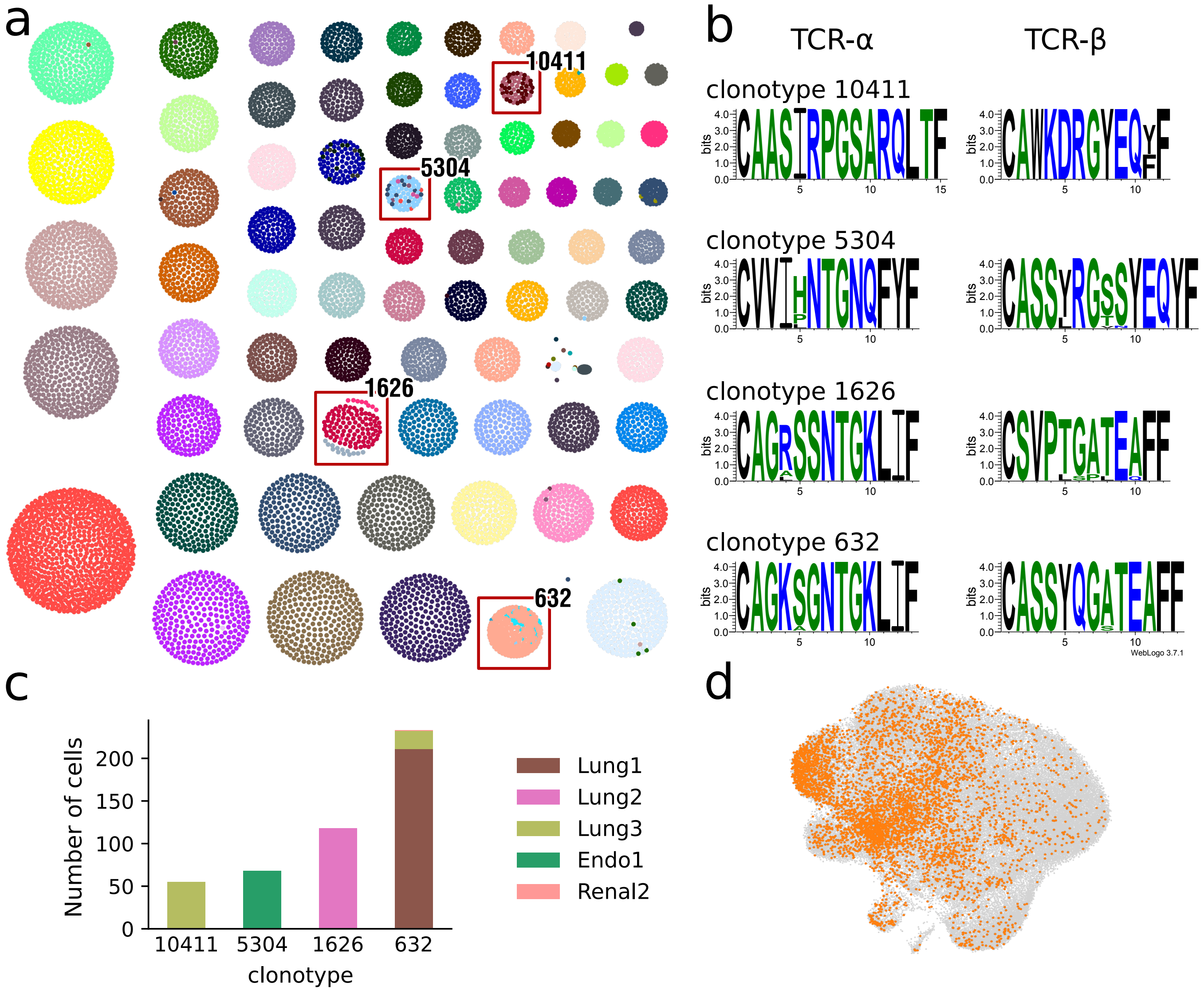


**Supplementary Figure 4: The comparison of clonotypes based on either sequence similarity or identity yields evidence of convergent evolution.** **(a)** Clonotypes with more than 50 cells visualized as a network plot. Each cluster represents a sub-network corresponding to a clonotype based on amino-acid sequence similarity; each dot represents an individual cell and is colored according to the clonotype assigned based on nucleotide-sequence identity. Four clonotypes with heterogeneous nucleotide sequences are highlighted. **(b)** Sequence logos of the primary TCR-α and TCR-β chains of the clonotypes highlighted in (a). **(c)** Patient composition of the four clonotypes highlighted in (a). Clonotypes number 10411, 5304, and 1626 comprised cells from a single patient only and were likely the result of convergent clonotype evolution. Clonotype 632 comprised cells from different patients and potentially represents an epitope targeting a common disease epitope (e.g. a viral epitope). **(d)** Cells from convergent clonotypes highlighted in the UMAP plot from Wu *et al.* [1]. We defined convergent clonotypes as subnetworks including different nucleotide sequences from the same patient.

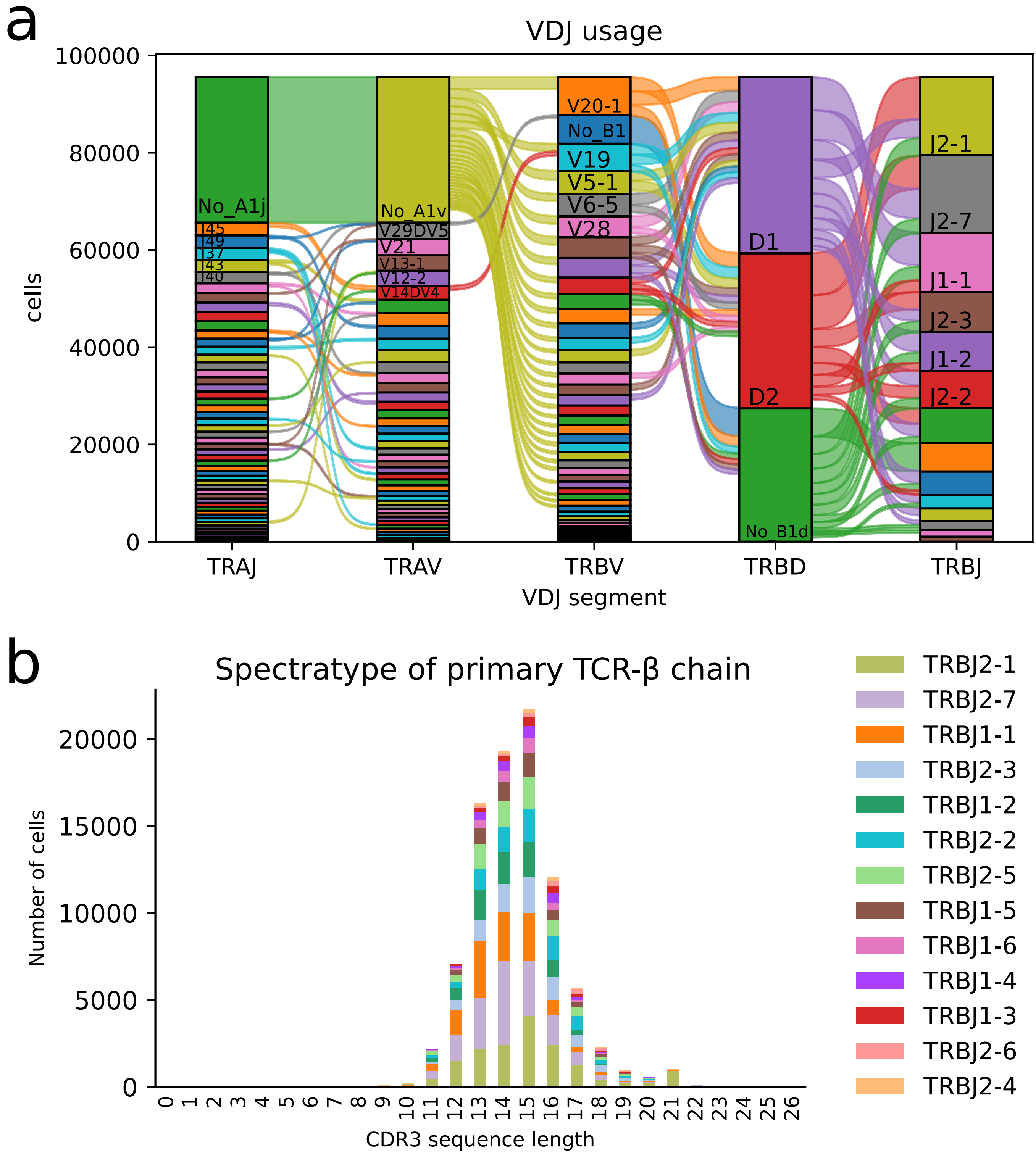


**Supplementary Figure 5: VDJ gene usage of 96,000 T cells from Wu *et al.*** [1] **(a)** VDJ gene usage of primary TCR-α and β chains visualized as a Sankey plot. Each bar refers to a specific VDJ segment, and each bar section to a specific V, D, or J gene. **(b)** Spectratype plot of the primary TCR-β chain, colored by J gene usage. The bars show the distribution of TCR-β CDR3 sequence length.

| Tool name | Website | Language | Receptor type | Support for 10x | Support for Smart-seq2 | Paired chains | Post-processing of clonotypes | Basic IR visualization | Advanced IR visualization | Clonotype clusters of single cells | Integrated with Gex | Ref. |
|---|---|---|---|---|---|---|---|---|---|---|---|---|
| Immcantation (pRESTO) | https://immcantation.readthedocs.io | Python (and R) | TCR BCR | ✗ | ✗ | ✗ | ✓ | ✓ | ✓ | ✗ | ✗ | [3] |
| ImmuneArch (tcR) | https://immunarch.com/ | R | TCR BCR | ✓ | ✗ | ✗ | ✓ | ✓ | ✓ | ✗ | ✗ | [4] |
| iMonitor | https://github.com/zhangwei2015/IMonitor | Perl (and R) | TCR BCR | ✗ | ✗ | ✗ | ✓ | ✓ | ✗ | ✗ | ✗ | [5] |
| VDJtools | https://github.com/mikessh/vdjtools | Java (Python) | TCR BCR | ✗ | ✗ | ✗ | ✓ | ✓ | ✓ | ✗ | ✗ | [6] |
| TRUST4 | https://github.com/liulab-dfci/TRUST4 | Perl | TCR | (✓)[1] | (✓)[1] | ✗ | ✗ | ✗ | ✗ | ✗ | ✗ | [7] |
| scTCR Seq | https://github.com/ElementoLab/scTCRseq | Python | TCR | ✗ | ✓ | ✓ | ✓ | ✓ | ✗ | ✗ | ✗ | [8] |
| TraCeR | https://github.com/teichlab/tracer | Python | TCR | ✗ | ✓ | ✓ | ✓ | ✓ | ✗ | ✓ | (✓)[2] | [9] |
| VDJ Puzzle | https://github.com/simone-rizzetto/VDJPuzzle | Java | TCR | ✗ | ✓ | ✗ | ✓ | ✓ | ✗ | ✗ | (✓)[2] | [10] |
| TRAPes | https://github.com/YosefLab/TRAPeS | C++ | TCR | ✗ | ✓ | ✗ | ✗ | ✗ | ✗ | ✗ | ✗ | [11] |
| Scirpy | https://github.com/icbi-lab/scirpy | Python | TCR | ✓ | ✓ | ✓ | ✓ | ✓ | ✓ | ✓ | ✓ | |

**Supplementary Table 1:** Comparison of Scirpy with currently available tools supporting immune repertoire analysis at the single-cell level, or offering immune repertoire visualization. Basic immune repertoire (IR) visualization includes clonotype abundance, diversity, and VDJ usage. Advanced IR visualization also includes repertoire overlap, clustering as well as specialized analysis of individual clonotypes and publication ready figures. "Gex" indicates gene expression data.

[1] Requires extra settings.
[2] Requires special skills and extra coding.

### 1. Supplementary Note

A clonotype designates a collection of T or B cells that bear the same adaptive immune receptors, and thus recognize the same epitopes. Generally, these cells are also descendants of a common, antecedent cell and belong to the same cell clone. In single-cell RNA-sequencing (scRNA-seq) data, T cells sharing identical complementarity-determining regions 3 (CDR3) sequences of both α and β TCR chains make up a clonotype.

Contrary to what would be expected based on the previously described mechanism of allelic exclusion [12], scRNA-seq datasets can feature a considerable number of cells with more than one TCR α and β pair. Since cells with more than one productive CDR3 sequence for each chain did not fit into our understanding of T cell biology, most TCR analysis tools ignore these cells [13, 14] or select the CDR3 sequence with the highest expression level [11]. While in some cases these double-TCR cells might represent artifacts (e.g. doublets of a CD8+ and a CD4+ T cell engaged in an immunological synapse), there is an increasing amount of evidence in support of a *bone fide* dual-TCR population [15, 16].

Scirpy allows investigating the composition and phenotypes of both single- and dual-TCR T cells by leveraging a T cell model similar to the one proposed in [9], where T cells are allowed to have a primary and a secondary pair of α and β chains. For each cell, the primary pair consists of the α- and β-chain with the highest read count. Likewise, the secondary pair is the pair of α/β-chains with the second highest expression level. Based on the assumption that each cell has only two copies of the underlying chromosome set, if more than two variants of a chain are recovered for the same cell, the excess TCR chains are ignored by Scirpy and the corresponding cells flagged as "multichain" (Supplementary Figure 1). This filtering strategy leaves the choice of discarding or including multichain cells in downstream analyses.

Scirpy implements a network-based clonotype definition that enables clustering cells into clonotypes based on the following options:

(a) identical CDR3 nucleotide sequences;

(b) identical CDR3 amino acid sequences;

(c) similar CDR3 amino acid sequences based on pairwise sequence alignment.

The latter approach is inspired by studies showing that similar TCR sequences also share epitope targets [13, 17, 18]. Based on these approaches, Scirpy constructs a global, epitope-focused cell similarity network. While convergence of the nucleotide-based clonotype definition to the amino acid based one hints at selection pressure, sequence alignment based networks offer the opportunity to identify cells that might recognize the same epitopes.
