## Supplementary figures and images for "Scirpy: A Scanpy extension for analyzing single-cell T-cell receptor sequencing data"

### figure1 (high resolution)

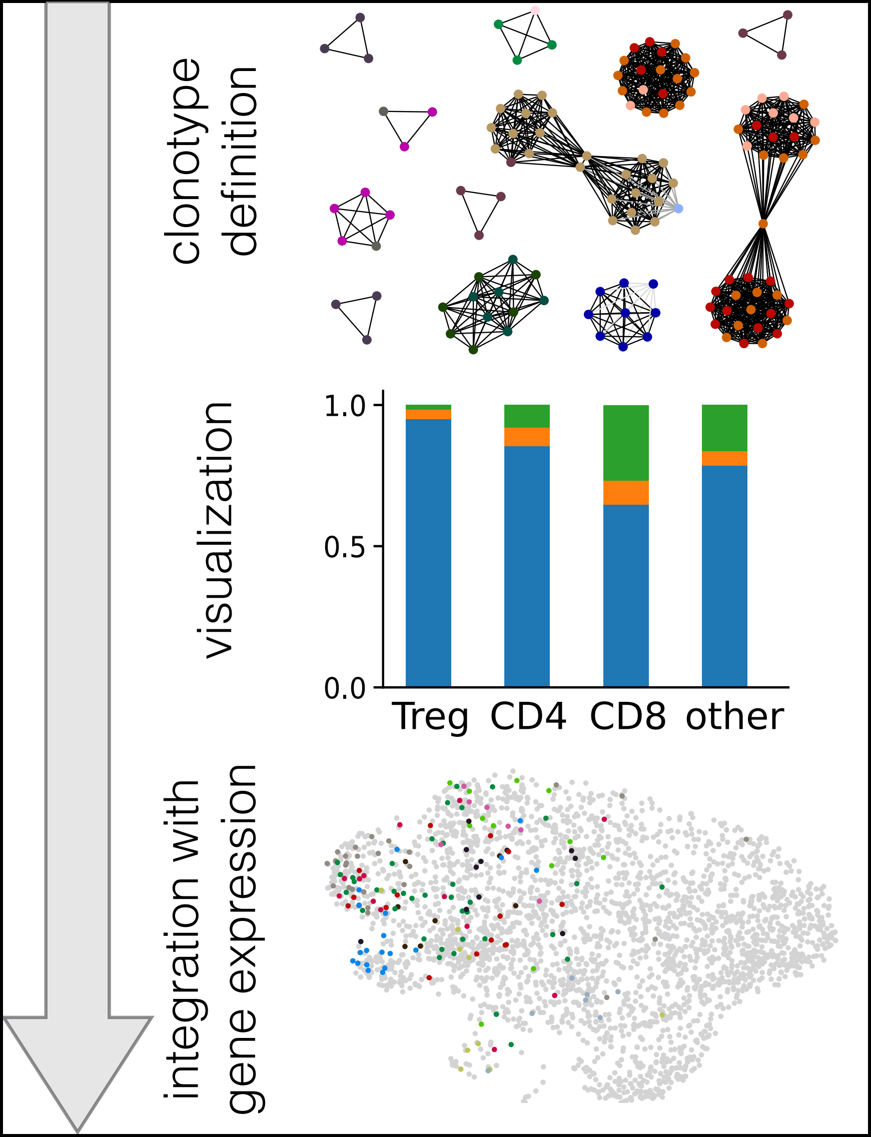
